# Active surveillance for influenza A virus in swine reveals within-farm reassortment and cocirculation of distinct subtypes and genetic clades

**DOI:** 10.1101/2024.06.28.601245

**Authors:** Megan N. Thomas, Garrett M. Janzen, Alexey Markin, Aditi Sharma, Kelly Hewitt, Ganwu Li, Amy L. Baker, Phillip C. Gauger, Tavis K. Anderson

## Abstract

Influenza A virus (IAV) is one of the three most frequently detected respiratory pathogens in swine. A passive IAV in swine surveillance system in the United States (U.S.) provides aggregated national metrics to quantify spatial and temporal changes in genetic diversity.

However, swine production is not homogenous: production systems vary in size and management strategies that affect the transmission and evolution of IAV. To assess the impact of fine-scale variation in swine production on IAV transmission, we conducted active surveillance on sow farms and linked nurseries from 4 U.S. production systems for up to 14 monthly collections. From IAV-positive samples, we obtained 85 complete HA sequences, and of these, we successfully assembled 62 whole genomes with associated epidemiological information. To infer transmission and evolution, we conducted Bayesian phylodynamic analyses and detected six genetic clades from four HA lineages: the H1 1A classical swine, the H1 1B human-seasonal, and the H3 2010.1 and 1990.4 lineages. The 1B and H3 1990.4 strains showed evidence of transmission from sow farm to nursery. In contrast, 1A and H3 2010.1 viruses were detected in nurseries without detection in a linked sow farm. We also detected seven separate human-to-swine transmission events in the H1N1 pandemic clade (1A.3.3.2) in sow and nursery sites. Our data demonstrated that nursery sites were infected with IAV that was both linked and unlinked to the sow farm. These data suggest that control efforts may be impacted by subclinical IAV transmission in the breeding herd, the mixing of sow farm sources at the nursery, regional spread of new strains, and human-to-swine transmission. Regular surveillance activities within production systems provide the ability to match vaccine components to circulating diversity, thereby minimizing the opportunity for novel reassorted viruses to emerge and impact animal health.

## Introduction

Influenza A viruses (IAV) are one of the most important respiratory pathogens of swine, often leading to high morbidity during outbreaks in swine herds [1]. The current genetic diversity of IAV in swine has been shaped by interspecies transmission of IAV from non-swine hosts. In the United States, all observed endemic IAV in swine lineages are derived from human seasonal IAV [2]. Following interspecies transmission, the RNA genome allows for evolution through mutation, particularly in the hemagglutinin (HA) and neuraminidase (NA) surface glycoproteins, or through reassortment, wherein gene segments are exchanged during viral replication in cells infected with two or more strains [3]. These processes can facilitate immune evasion through antigenic drift and shift [4–6] and facilitate the adaptation of human IAV to the new swine host [7,8]. The quintessential example of this dynamic is the 2009 H1N1 pandemic (1A.3.3.2) lineage: this virus became established in swine following human-to-swine spillover, evolved and emerged as the 1A.3.3.2 clade following reassortment in swine, and has subsequently been consistently reintroduced to pigs [9,10]. A direct consequence of IAV evolution and interspecies transmission of IAV has been that vaccine control in pigs is complex due to an array of virus genotypes and antigenic profiles.

Vaccines used in swine often do not completely block infection under typical field situations [11,12]. Passive or previously acquired immunity against genetically similar viruses can protect pigs against clinical signs but not against infection or transmission [13,14].

Additionally, prior immunity from infection, vaccination, or maternal antibodies can shape the immunological profile of animals that may interfere with vaccine efficacy, the active immune response to vaccines, and the likelihood that an IAV infects and transmits within herds [15–17]. Phylogenetic analyses with national surveillance data have demonstrated at least 15 clades across the H1N1, H1N2, and H3N2 subtypes detected within the last two years with regional differences [18–20], and reassortment between endemic viruses appears to alter the evolutionary dynamics of the surface proteins [21,22]. In attempts to control this diversity, vaccine platforms have been developed to incorporate custom antigens [23], obviate the impact of maternal antibodies [24,25], or pivot to incorporate antigens detected in on-farm surveillance. While all of these approaches prove effective in controlled experimental systems, modern swine agriculture has unique abiotic and biotic conditions that change IAV transmission, and surveillance detections in the United States demonstrate year-round circulation of IAV [18,26]. Intensified swine production systems and interstate movement of pigs in the United States likely enables the introduction, spread, and maintenance of IAV in different age classes within and between swine production systems [21,27,28].

Swine farms in North America generally use a multi-site production process, allowing for the optimization of biosecurity processes and environmental specifications for the developmental stages of the pigs. Breeding stock have the highest economic value, represent significant investment in genetic improvement, and are housed at “nucleus sites.” Consequently, these sites have the highest levels of biosecurity, but animals may be moved to multiple locations and have the potential to carry IAV into “downstream” production stages. Sow farms house selected breeding females for multiple reproductive cycles (∼20 weeks of pregnancy, lactation, and weaning) and have either been vaccinated against or naturally infected with IAV, commonly by more than one strain, and have a dynamic immune response to IAV exposure [12,29]. In contrast, neonatal piglets on the sow farm are born immunologically naïve but may be protected from infection based on the degree by which they aquired maternally derived antibodies [30,31]. Given the multitude of factors contributing to the transfer of passive immunity, levels of IAV detection lon sow farms exhibit significant variation and can persist endemically, even at very low levels of circulation [32–34]. In some production models, weaned piglets are moved to a nursery facility and later to a finisher site, growing to market-ready weights of 280-300 lbs.

These wean-to-finish and finisher sites are the final stage of production, have the lowest levels of biosecurity, and have been associated with IAV outbreaks, in which a high proportion of the pigs become clinically ill in a relatively short period [35]. Consequently, the multi-site swine production model that is economically favorable has created a unique opportunity for pathogen evolution: maternal antibodies and prior infection or vaccination create a heterogeneous immune landscape; animal movement introduces naïve or infected hosts to new locations; and infection and reassortment may lead to the emergence of novel IAV that makes targeted vaccine control challenging.

In this study, we conducted active surveillance for IAV in four swine production systems, using one sow farm and the connected nurseries as a proxy for the system. Our goal was to determine the major transmission pathways for IAV through fine-scale farm-level surveillance.

To identify modes of IAV transmission and management-associated risk factors for IAV detection, we used Bayesian phylogenetic and network modeling techniques. With these methods, we demonstrated direct sow farm-to-nursery transmission, highlighted how production system management influences farm-level IAV risk, and reinforced the importance of the human-swine interface for IAV transmission to pigs. This study demonstrated significant IAV diversity within production systems, and evolutionary dynamics and production practices within a swine system that can be used to inform farm-level management.

## Methods

### Farm enrollment and characterization

Four sow farm and nursery combinations, referred to as Farms A-D, were selected as a representative of their respective production system based on the following criteria: evidence of endemic IAV circulation based on previous diagnostic submission to the Iowa State University Veterinary Diagnostic Laboratory (ISU-VDL), representation of modern swine production practices and facilities, and different geographic distribution. The four farms include sow and nursery sites in Missouri, Iowa, Nebraska, South Dakota, Minnesota, and Oklahoma. Farm B was a high-value breeding stock herd (genetic nucleus), and Farms A, C, and D were commercial production herds. The sampled sow farm, with one exception (described below), remained the same throughout the study, while the sampled nursery changed based on the schedule, which determines where the piglets from the sow farm were placed. The veterinarian overseeing each farm completed a 15-question survey to acquire information regarding the site’s structure, production and health parameters, influenza vaccination history, and biosecurity practices (Supplemental File 1). Each farm used autogenous or custom IAV vaccines, with the HA gene sequences corresponding to the farm’s vaccine antigens being provided.

### Sample collection supplies and procedures

Active sampling was conducted at each farm (sow and linked nursery) once a month from May 2020 to June 2021. At the sow farm, udder wipes from the sow (UW; n=10) and nasal wipes from the piglets (NW; n=16) were collected when the suckling pigs were 2- to 3 weeks of age. At the nursery, nasal wipes (n=16) and oral fluids (OF; n=4) were collected from the group of weaned pigs sampled at the sow farm the previous month, now 6 to 9 weeks of age. Samples (n=26 from sow farm, n=20 from nursery) were chilled before transport to the ISU-VDL for subsequent analyses. Deviations from this process included Farm A, which, in July 2020, began to limit the handling of suckling piglets and could no longer provide nasal wipes. Instead, Farm A provided 15 to 26 udder wipes for the remainder of the study. Farm B experienced a disease outbreak in the sow farm in December 2020, leading to the discontinuation of sampling udder and nasal wipes from this site. When sampling was re-initiated in March 2021, the samples were taken from a different sow farm within the same production system. Farms A and B participated in 13 months of sampling, while Farms C and D participated in 9 months of sampling, defined by submission of at least one set of samples (sow farm or nursery).

Udder wipe kits, nasal wipe kits, and oral fluid kits were prepared at the ISU VDL and shipped to a location in the production system that distributed the sample collection supplies to the sow farm and appropriate nursery location. Udder wipe and nasal wipe kits consisted of 10 mL of Dulbecco’s modified eagle medium (DMEM, Thermo Fisher Scientific, Waltham, MA, USA) placed in a Whirl-Pak® 710 mL bag (Sigma-Aldrich, Inc., St. Louis, MO, USA) and 1 Fisherbrand 4” x 4” sterile gauze (Fisher Scientific, Waltham, MA, USA). The gauze was moistened with DMEM and wiped across the udder skin of the dam or the nose of a pig [36,37]. The gauze was placed into the whirl-pak bag, secured to prevent leakage, and submitted to the ISU VDL. Oral fluid kits consisted of 1 zip-lock bag, 50 mL conical tube, and 3-4 feet of cotton rope (WegRigging Supply, Barrington, IL, USA). Participating farms were instructed to collect oral fluids by hanging the rope in the pen to shoulder height of the nursery pigs for 30 minutes, squeezing the rope in the zip-lock bag to harvest the fluid, and transferring the fluid to the 50 mL conical tube [38]. Oral fluids were submitted to the ISU VDL for diagnostic testing.

### Sample processing, RNA extraction, reverse transcription real-time polymerase chain reaction, and sequencing

Porcine udder wipes, nasal wipes, and oral fluid samples collected at swine farms were shipped to the ISU VDL. After arrival, approximately 3-4 mL of udder and nasal wipe media or oral fluids were transferred to 5 mL tubes for storage or diagnostic testing. Before extraction, udder and nasal wipes were pooled in groups of 5 using 100 μL of each sample. Oral fluids were tested individually.

IAV extraction and reverse transcription real-time PCR (RT-rtPCR) were performed at the ISU VDL according to the standard operating procedures (SOP). Viral RNA extraction was performed using 100 μL of the sample and a MagMAX™ pathogen RNA/DNA isolation kit (Thermo Fisher Scientific, Waltham, MA, USA), Kingfisher96 instrument (Thermo Fisher Scientific, Waltham, MA, USA) per the manufacturer’s instruction. IAV RT-rtPCR was performed on nucleic acid extracts at the ISU VDL according to the SOP using the VetMAX™ Gold SIV detection kit (Thermo Fisher Scientific, Waltham, MA, USA) according to the manufacturer’s instructions. The RT-rtPCR was performed using standard mode on an AB 7500 Fast thermocycler (Thermo Fisher Scientific, Waltham, MA, USA). HA and NA subtyping was performed on RT-rtPCR–positive IAV nucleic acid extracts at the ISU VDL according to the SOP using the VetMAX™ Gold SIV subtyping kit (Thermo Fisher Scientific, Waltham, MA, USA) per manufacturer’s instructions. Samples with Ct <38 were considered positive for both detection and subtyping PCR.

For sequencing, viral RNA was extracted from infected udder wipes, nasal wipes, or oral fluids as previously described [39]. Next-generation sequencing was performed on an Illumina, Inc., MiSeq platform by following standard Illumina protocols at the ISU VDL [39,40]. Bioinformatics analyses to assemble each segment sequence of swine influenza virus were performed as described previously [41].

### Quantifying genetic diversity and inferring IAV transmission routes

We constructed a dataset of sequences representing IAV detected in the general swine population to complement our active surveillance sequence data. The general swine population sequences served as a genomic epidemiological control for our ancestral state reconstruction and allowed the detection of within-system transmission dynamics. The “Control” sequences were identified by querying the ISU VDL database for viruses with the same HA genetic clade as those sampled in the study. Inclusion criteria included collection from a state represented in the study and independence from the production systems to which the four study farms belong.

Additionally, we included human sequences (seasonal H3N2 and 2009 H1N1 pandemic) in our analysis by representatively sampling 20 H3N2 sequences from 17,667 sequences collected between 1990 and 2022 and 50 H1N1 sequences from 512 sequences collected between 2019 and 2021 using PARNAS to preserve the representation of genetic diversity [42].

To determine transmission routes, we applied phylodynamic analyses and identified an epidemiological linkage between sow and nursery locations. The phylogenetic pattern of sow farm-to-nursery transmission appears within a phylogeny as a nursery detection IAV gene sequence nested within a group of sow farm IAV gene sequences (denoted by color). The HA sequences were divided into subsets by lineage (H1-1A, H1-1B, and H3), aligned separately with mafft v7.490 [43], and a maximum-likelihood phylogenetic tree was inferred using FastTree v2.1.11 [44] as the input gene tree for subsequent analyses. Ancestral state reconstruction was performed on each lineage dataset using TreeTime v0.8.6 with farm site (Farm A-D and Sow Farm or Nursery) as the discrete trait [45]. The alignment and maximum-likelihood phylogenetic trees were used as input, and default parameters were retained. Ancestral state reconstruction was also performed using a Bayesian approach with BEAST v1.8.4 [46], where the temporal signal in the dataset was assessed using TempEst v1.5.3 [47]. The Bayesian analysis implemented a GTR substitution model with gamma-distributed rate variation, an uncorrelated relaxed clock [48], and the GMRF Skyride population demographic model [49]. Inferred ancestral relationships were visualized on a maximum clade credibility (MCC) tree, generated using TreeAnnotator [50], and visualized and colored by inferred ancestral state using FigTree [51].

### Detecting IAV reassortment within swine production systems

We assembled reference datasets of swine H1 and H3 whole genomes from the USDA IAV in swine surveillance system. We downloaded n=76,629 swine IAV gene sequences submitted by USDA surveillance from the NCBI Virus database [accessed December 2, 2023] [52]. We maintained only those strains with complete sequences for each of the eight gene segments, resulting in n=1,763 H1 and n=1,100 H3 whole genomes. All maintained gene segments were classified into clades according to global swine IAV nomenclature [53] using octoFLU [54]. Next, we assembled whole genomes associated with each HA gene collected in this study. Since many collected samples contained multiple alternative sequences for some gene segments, we established whole genome linkage by examining similar WGS strains in the reference dataset. This approach established a genetic similarity threshold based on the assumption of preferential pairing [21,22], i.e., for a sample with 2 NS genes, this approach determined the most likely NS gene to include within the whole genome. For each query HA collected in this study, we identified a set of *relevant strains* from the reference dataset. We called a reference strain *relevant* if its HA was within the 3% sequence divergence threshold from the query HA: this threshold was sufficient to find at least one reference strain for each query HA. Then, if there were multiple non-HA genes associated with the query HA, we linked the query HA with the gene that was most similar to one of the relevant strains. Similarly, if, for a non-HA segment, no gene sequence was obtained from a sample, we filled in this gap by considering the closest relevant reference strain and borrowing the segment sequence from it. This approach ensured that we minimized the number of artificiallyintroduced reassortment events.

We merged the reference data with the assembled whole genomes and inferred phylogenetic trees for each gene segment with separate phylogenies for H1 and H3 HAs using FastTree v2.1.11 [44]. To identify reassortment events within the production systems, we applied TreeSort v.0.1.1 (https://github.com/flu-crew/TreeSort) under the default parameters.

### Estimating farm-level IAV status with a predictive Bayesian network model

To predict whether a sample collected within a farm would test positive or negative for IAV, we developed a statistical framework employing Bayesian networks (BN) using the bnlearn R package [55]. Bayesian networks are directed acyclic graphs (DAGs) that link variables based on conditional probabilities. Each node in the graph represents a variable, and the links between each variable represent dependencies/causalities. The DAG represents the structure of the probability distribution, and we implemented it to predict the probability a pooled sample would test positive or negative for IAV, conditional on the predictor variables that described farm management and prior IAV detection results.

Due to management differences, separate models were built for samples from sow farms and nurseries. The unit of the response variable was positive or negative IAV detection for a pooled sample. Logically required edges were whitelisted, and logically inconsistent edges were blacklisted (Supplemental File 2). To determine the effect of past IAV-positive and IAV-negative pooled samples on the conditional probability of positive detection for a sample, we generated a categorical lag effect. The lag encodes whether a positive case was detected in the previous month in a sow farm or nursery from the same production system or in the same premise as the sample in question. The dataset contained missing data due to incomplete participation during the study and the lack of records of IAV cases in the temporal lag variables in the first month.

For this reason, we used Structural Expectation-Maximization (Structural EM) [56], a method designed to accommodate missing data in BN structure learning. The maximization step of the learning algorithm was carried out using the hill-climbing greedy search procedure. To investigate the role of the predictor variables in determining IAV status, conditional probabilities were queried using the cpquery function in R using probabilistic logic sampling [57] with 10,000,000 draws per inference.

## Results

### Management practices varied across the sampled swine production systems

Management practices differed among the four farms (Table 1). The nursery facilities varied in the degree of mixing of weaned piglets from different sow farms: Farms B and C mixed rooms, barns, and sites, Farm D mixed only site and barn, and Farm A allowed no mixing at any level. The number of sows (females with at least one litter) in the sow farms varied from 700 to 7,600, with the percentage of gilts (females that have not yet had a litter) ranging from 20 to 60%. All farms except Farm B purchased gilts externally, while Farm B replaced gilts from within the system. In all four farms, gilts received two doses of autogenous IAV vaccine during the development phase, generally from 22 to 30 weeks of age. For the gilts, age at vaccination, interval between doses, and vaccine antigens differed between farms. All farms except Farm D practiced biannual sow herd vaccination with an autogenous IAV vaccine.

**Table 1.**
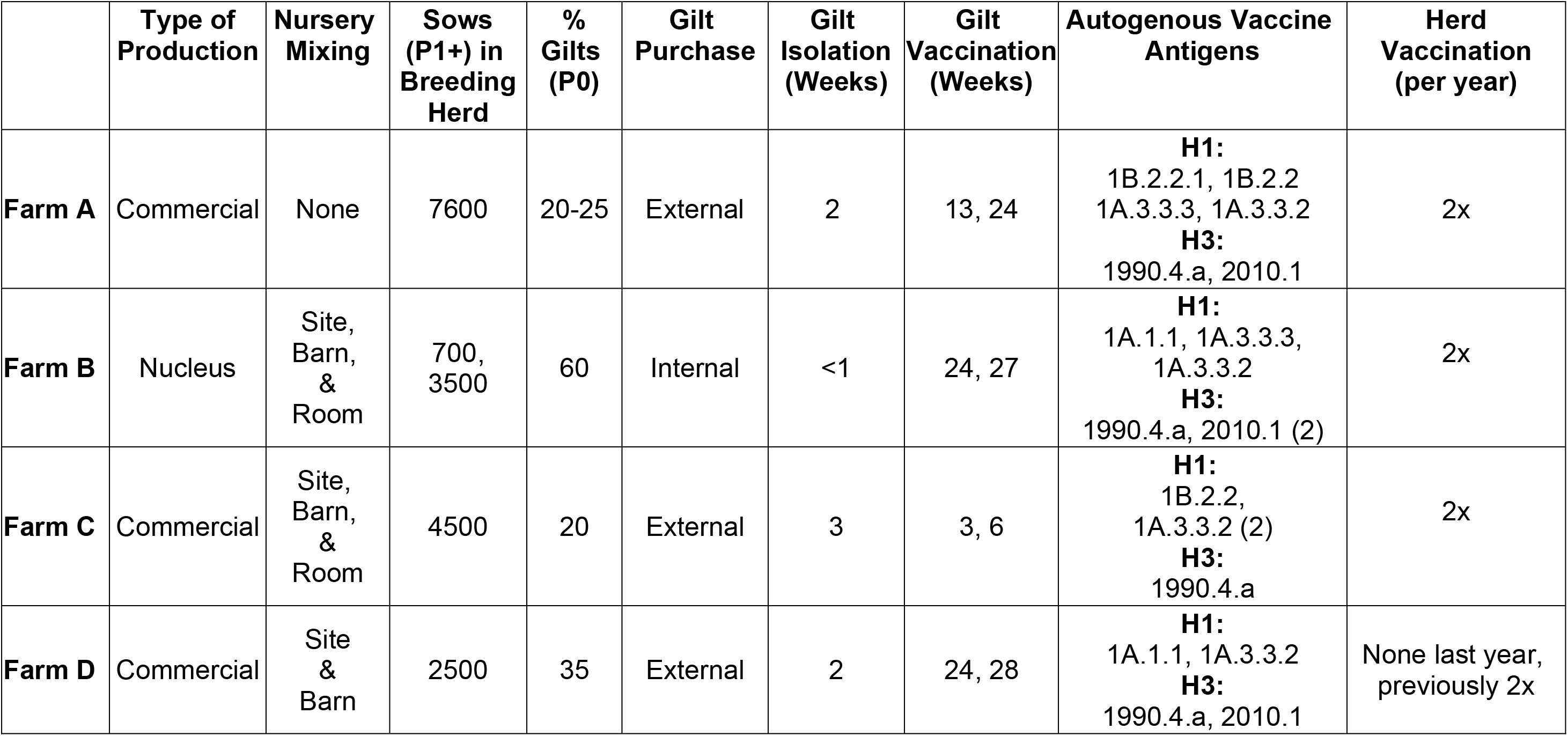
Results of production practices survey administered to farm veterinarians.

### Influenza A virus detection and prevalence

Farms A, B, and C shared a general pattern in which IAV was detected somewhere in the system in nearly all sampled months, and their respective nurseries had more positive months than their sow farms (Figure 2). Specifically, Farm A was positive at least once within the system for 93% (13/14) of months sampled, with the sow farm positive 46% (6/13) of months and the nurseries positive 77% (10/13) of months. The Farm A system had no sow farm submission in January and no nursery submission in December. Farm B was positive 92% (12/13) of months, with the sow farm positive 22% (2/9) of months and the nurseries positive 85% (11/13) of months. Farm C was positive 91% (10/11) of months, with the sow farm positive 56% (5/9) of months and the nurseries positive 100% (9/9) of months. Farm D had the lowest participation level, and within these data, IAV was detected in the system in only 50% (5/10) of months, 50% (3/6) of months in the sow farm and 44% (4/9) months in the nurseries. The sample positivity (PCR positive samples divided by all samples) for each Farm A-D followed a similar pattern, with nurseries being having more positives than the sow farm. Farm A had an overall positivity of 56% (80/143 samples), a sow farm positivity of 44% (27/61), and a nursery positivity of 65% (53/82). Farm B had an overall positivity of 39% (52/133), a sow farm positivity of 16% (7/45), and a nursery positivity of 51% (45/88). Farm C had an overall positivity of 61% (63/104), a sow farm positivity of 42% (19/45), and a nursery positivity of 75% (44/59). Lastly, Farm D had an overall positivity of 33% (30/90), a sow farm positivity of 30% (9/30), and a nursery positivity of 35% (21/60).

**Figure 1.**
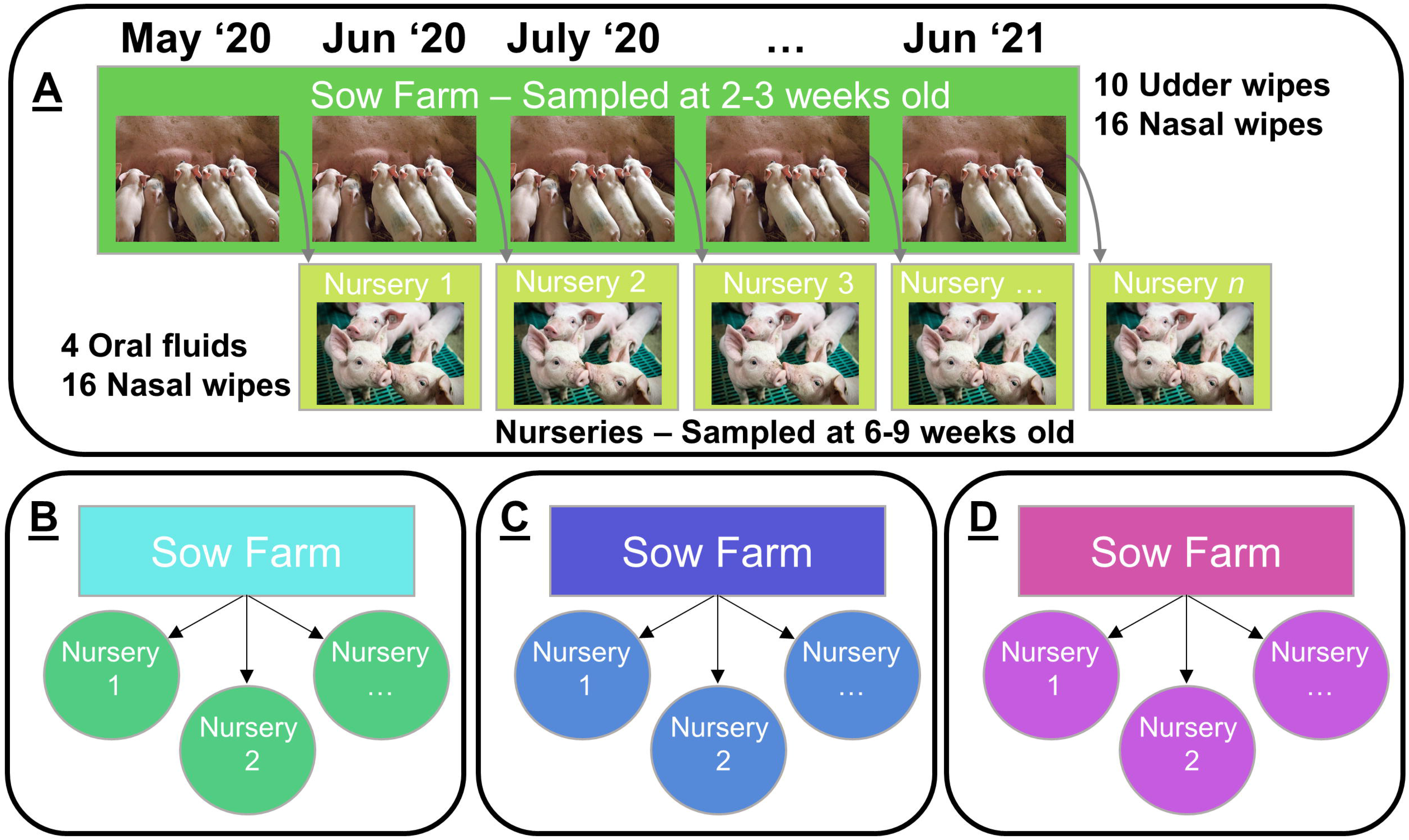
Active surveillance sample collection framework. Sampling began in May 2020 and continued until June 2021. Details of the sampling protocol are listed for Farm A and are identical for Farms B-D. For each Farm, one sow farm and multiple corresponding nurseries were sampled. Grey arrows in panel A represent movement of weaned pigs from the sow farms to the respective nursery. The number of nurseries varied between farms. Sow farms were generally sampled with 10 udder wipes and 16 nasal wipes when piglets were 2-3 weeks old. Nurseries were generally sampled with 4 oral fluids and 16 nasal wipes when piglets were 6-9 weeks old.

**Figure 2.**
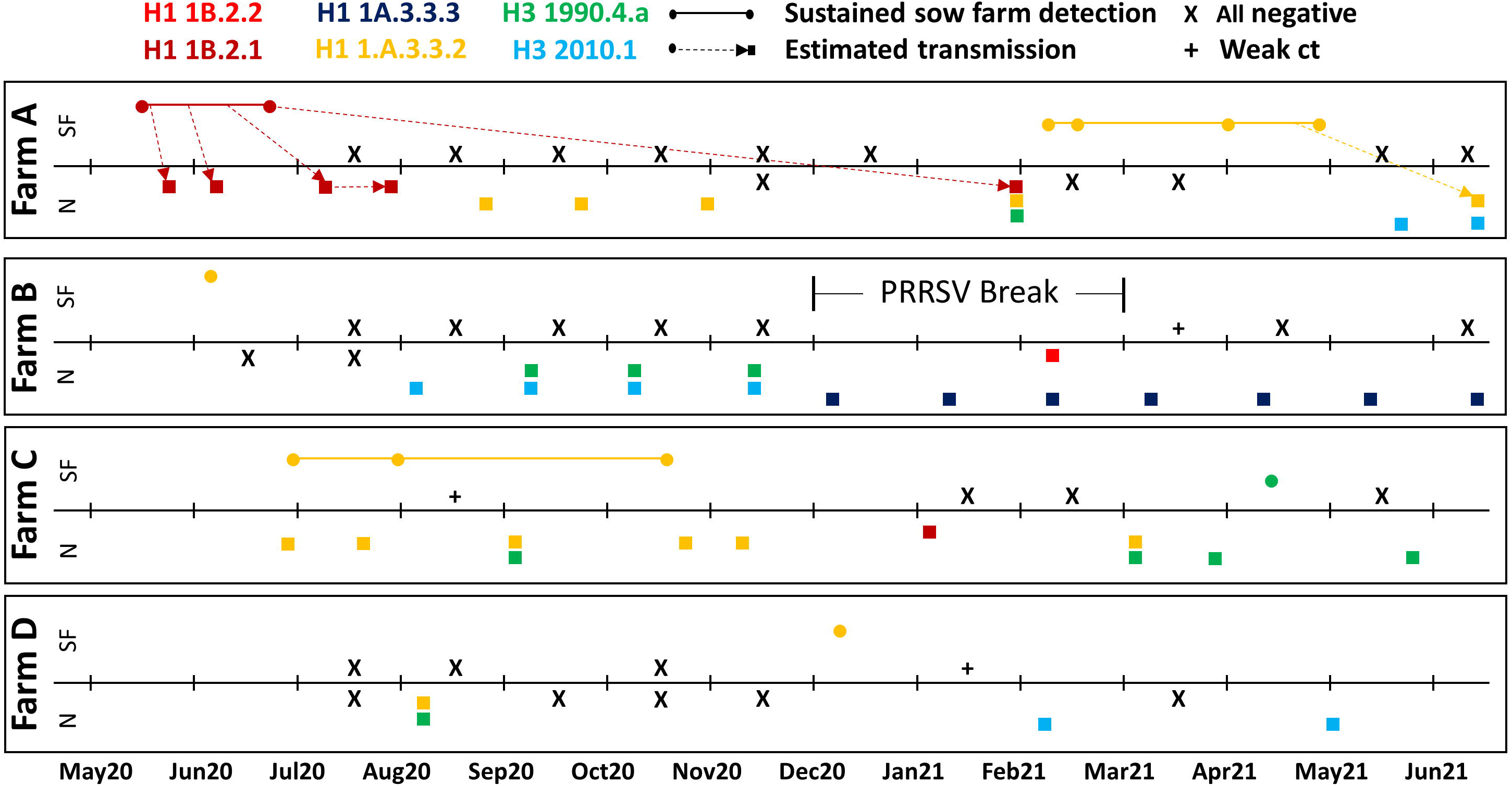
**Timeline of IAV sampling and detection in farms A-D from May 2020 to June 2021**. Circles represent sow farm sequences and squares represent nursery sequences. The circles and squares are colored by HA clade. Transmission between different locations was estimated using Bayesian phylodynamic techniques and is represented with a dashed line. A negative sampling event, depicted by a black X, indicated that all samples received for that month tested PCR negative for IAV. A sampling event with a weak Ct is depicted by a black plus sign. A “weak Ct” designation indicated that at least one sample from that date was PCR positive, but the amount of virus in the sample was too low for accurate sequencing. A pause in sample collection from the sow farm in Farm B is depicted with the text “PRRSV Break” between two brackets.

Generally, there was concordance between sampling type and IAV detection in the sow farm and nursery. In 89% of complete sow farm sample collections (24/27), defined as including UW and NW, the IAV status was consistent. However, in one sow farm sampling time point (Farm D, January 2021), a single udder wipe pool was positive, while all nasal wipe pools were negative (no sequence recovered). Additionally, two other sow farm sampling time points (Farm C, July and August 2020) had a positive NW pool without concordance in any UW pools (a whole genome sequence was successfully recovered from one but not the other). In the nursery, there was concordance between sampling types in 93% (40/43) of complete sample collections, including NW and OF. Three nursery sampling time points (Farm A, May 2021, Farm B, May 2021, Farm C June 2020) had at least one positive OF without concordance in any NW pools (whole genome sequences were recovered from all three).

There were 225 IAV-positive sample pools out of the 470 sample pools collected between May 2020 and June 2021. Of the 225 positive pools, 62 were from a sow farm and 163 were from a nursery. At least one gene segment sequence was recovered from 19 sow farm and 60 nursery sample pools. For a subset of these sample pools (14/19 sow farm and 48/60 nursery), all eight gene segments were successfully sequenced and reconstructed into whole genomes. Of the sow farm sample pools that yielded all eight segments, 9 were NW, 3 were UW, 1 was an OF submitted outside of the study protocol by Farm A, and 1 sample type was not recorded. The average cycle threshold (ct) value from the detection PCR run on these fully sequenced sample pools was 28.2 (max 33.0). Regarding the nursery sample pools that yielded all eight segments, 19 were NW and 29 were OF. The average ct value from the detection PCR was 27.3 (max 32.9).

We reconstructed 62 whole genomes from 79 samples from which at least one segment was successfully sequenced (Supplemental Table 1). Multiple sequences of the same gene segment were recovered from the same sample pool. In total, 24 of the 79 sample pools had evidence of more than one sequence. The minimum evidence was the detection of multiple lineages of the same internal gene or multiple clades of HA/NA. Of these 24 samples, 9 had multiple HA genes, 12 had multiple NA genes, and 18 had multiple lineages of the same internal gene (PB2, PB1, PA, NP, M, or NS). Of the internal genes, multiple lineages were most commonly detected for the NS gene segment (n=9). The most frequently sequenced gene segments were non-structural protein 1 (NS1; n=134) and matrix (M1M2; n=121). Though not all identical, all 121 M1M2 genes were of H1N1pdm (pdm) lineage. The segments that were most limiting to recovering all eight gene segments into a complete genome sequence were the three polymerase genes (PA, PB1, and PB2; n=76, 75, 76).

### Farm-level hemagglutinin and neuraminidase diversity and temporal detection

All farms had H1 and H3 HA subtypes and N1 and N2 NA subtypes detected at least once during the 14-month study. An average of 3.75 distinct HA clades (min = 3, max = 5) and 3.25 distinct NA clades (min = 3, max = 4) were detected within a farm during this period (Figure 2). Six unique HA clades and five unique NA clades were detected. Half (3/6) of the unique HA clade detections in the sow farm were not detected again in the same farm in the subsequent months, while the other half were sustained for two to five months. Only one distinct HA and NA clade was detected at each sampling point on each sow farm. One to three (mean HA = 1.31, mean NA = 1.44) distinct HA or NA clades were detected at a single sampling point in the nurseries.

The most commonly detected HA clade was H1N1pdm (1A.3.3.2), accounting for 37.7% of HA sequences (32/85). The other HA clades detected in order of prevalence were H3 1990.4.a (20.0%; 17/85 sequences), H3 2010.1 (14.1%; 12/85 sequences), H1 1B.2.1 (14.1%; 12/85 sequences), H1 1A.3.3.3 (12.9%; 11/85 sequences), and H1 1B.2.2 (1.2%; 1/85 sequences). To compare these detection frequencies to national surveillance data, the most commonly detected HA clade from USDA IAV in swine surveillance data for the same six states over the same timeframe was H1 1A.3.3.3 (32.7%; 283/863 sequences), followed by H1 1B.2.1 (19.6%; 169/863 sequences) and H3 2010.1 (15.5%; 136/863 sequences) with H1 1A.3.3.2 representing 8.8% of detections (76/863). The 1A.3.3.2 clade was detected in every sow farm and three out of four sets of nurseries represented by the four participating farms, most commonly in Farm A (n = 14 sequences) and Farm C (n = 14 sequences).

The most commonly detected NA clade was N2 2002 at 41.94% of NA sequences (39/93). The other NA clades detected in order of prevalence were H1N1pdm (32.3%; 30/93), N2 1998 (12.9%; 12/93), N1 classical swine (11.8%; 11/93), and N2 LAIV-98 (1%; 1/93). The detection frequencies within national surveillance data for the NA clade were N2 2002 (36.9%; 306/829), followed by N1 classical swine (32.9%; 273/829), N2 1998 (20.3%; 169/829), H1N1pdm (8.1%; 67/829), and LAIV-98 (1.2%; 10/829).

### Evidence for different IAV transmission pathways and interspecies transmission

Ancestral state reconstruction of the 1A.3.3.2 HA clade revealed seven independent human-to-swine transmission events (Figure 4). Farm A had three independent spillover detections, Farm C had two, and Farms B and D had one each. The spillovers were estimated to persist within the respective farms for a minimum of 6 and a maximum of 24 months. They may have been detected for a more extended period if sampling had been continued. The spillovers were detected in both sow farms and nurseries. In one instance, there was evidence of the spillover strain spreading from the sow farm to a nursery (Figure 4).

**Figure 3.**
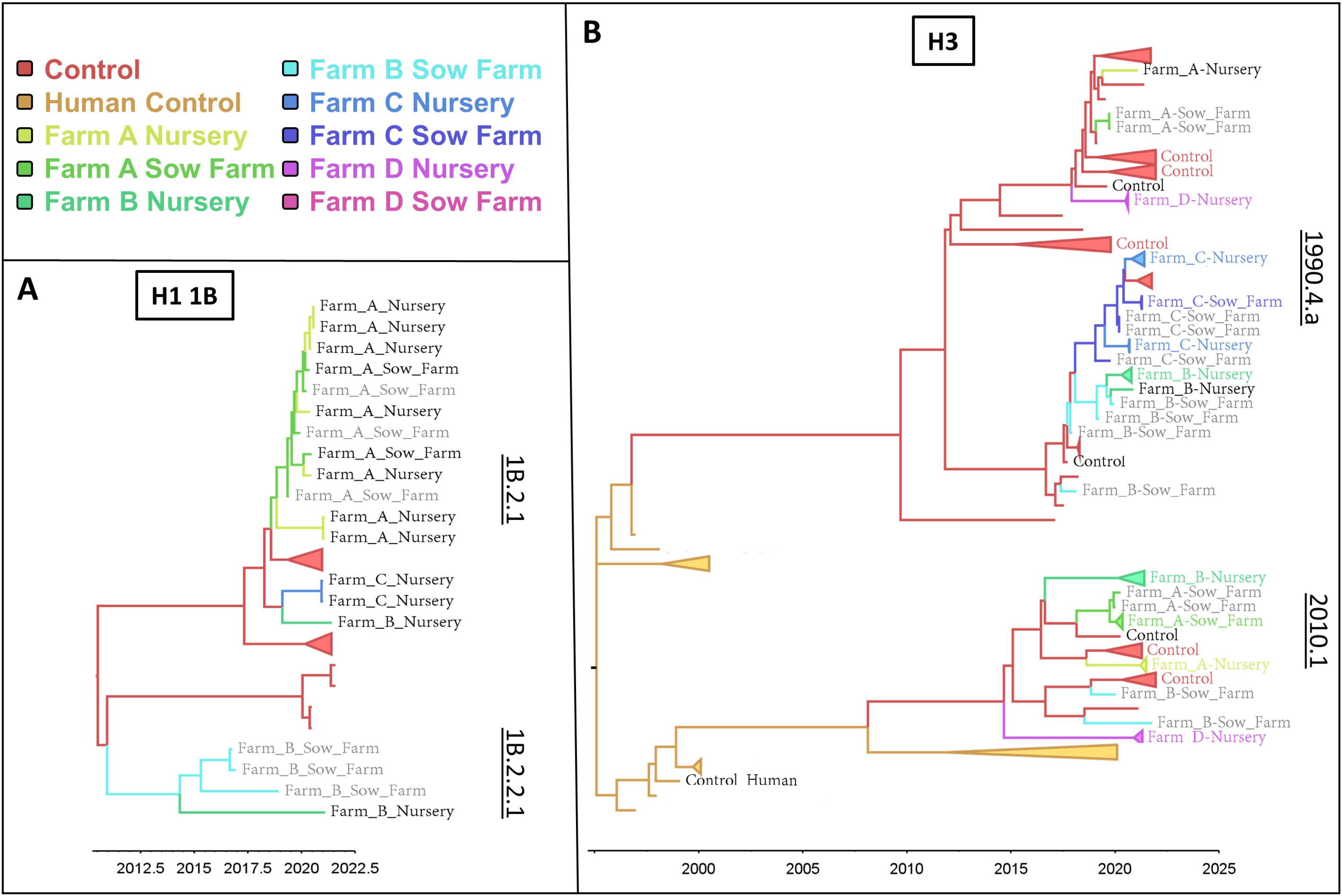
Time-scaled phylogenetic trees for H1 1B (A) and H3 (B) lineage influenza A virus. Tree branches are colored by inferred ancestral state of virus location: ancestral states were inferred using Bayesian phylodynamic techniques and consisted of discrete states. Tree tip labels are colored by sample location if resulting from a sampling event in this study. Tree tip labels in grey are historical sequences collected by the ISU VDL prior to the initiation of this study. The “Control” group contains influenza A virus sampled and sequenced from the general U.S. swine population outside of the four studied production systems. The “Human Control” group (B) contained human seasonal H3 IAV collected and sequenced in the U.S. between 1990- 2022. For easier visualization, some tree tips with shared sampling location are collapsed at the node and are represented by colored triangles.

**Figure 4.**
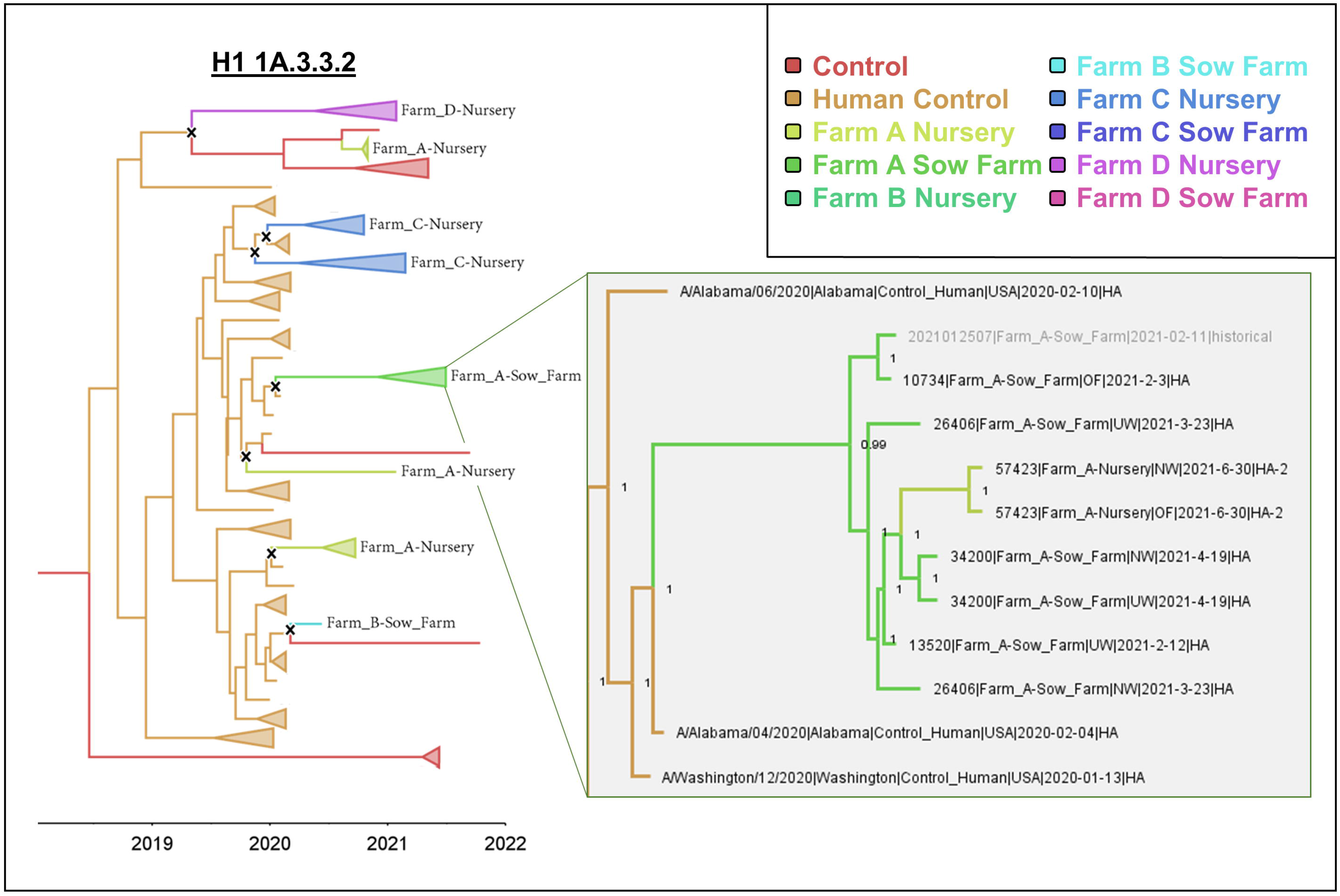
Time-scaled ancestral state estimation phylogenetic trees for H1N1pdm (1A.3.3.2) clade IAV. Tree branches and tips are colored as described in Figure 3. The “Human Control” group contains human seasonal H1N1pdm IAV in the U.S. from 2019 to 2021. An “X” denotes instances of human-to-swine spillover events. The inset portion of the tree depicts a human-to-swine spillover event into the sow farm of Farm A, which successfully transmitted to a nursery of Farm A.

On the phylogenetic tree, the pattern of sow farm-to-nursery transmission appeared as a nursery detection nested within a clade of associated sow farm detections (denoted by color).

Out of 33 IAV-positive nursery detections, 5 could be inferred as the result of sow farm-to- nursery transmission (Figure 3; Figure 4). When historical sow farm HA sequences were included alongside data generated in active surveillance, an additional nine nursery detections (total = 14/33) could be inferred from sow farm-to-nursery transmission. There were 19/33 detections in nurseries without any evidence (historical or otherwise) of a genetically related IAV in the sow farm. On the phylogenetic tree, these detections had a most recent common ancestor associated with the general U.S. swine population (a “control” sequence). There were no instances of inferred transmission from a nursery to a sow farm. When considering sow farm detections, 9/11 were associated with transmission to the nursery, and 2/11 were independent, without any evidence of transmission to a nursery.

### Evidence for reassortment within single production systems

We observed 15 putative reassortment events within the production systems. Many of these events were due to the introductions of novel H1N1pdm genes by human-to-swine spillovers. Two of four production systems had an active circulation of the 2019-20 seasonal H1N1pdm. For example, in Farm A, an H1-1A.3.3.3 strain obtained a new pdm NP gene from a 2019-20 human-to-swine spillover. A similar NP reassortment with a 2019-20 human seasonal pdm NP gene was observed on Farm B with the H1 1A.3.3.3 and H1 1.B.2.2.1 strains, the former affecting ten strains in total. In reverse, we observed reassortment of novel H1N1pdm strains with the swine-endemic genes. For example, Figure 5 depicts the fixation of a swine-enzootic TRIG NS gene in the clade of novel H1N1pdm viruses in Farm C that was maintained for at least eight months.

**Figure 5.**
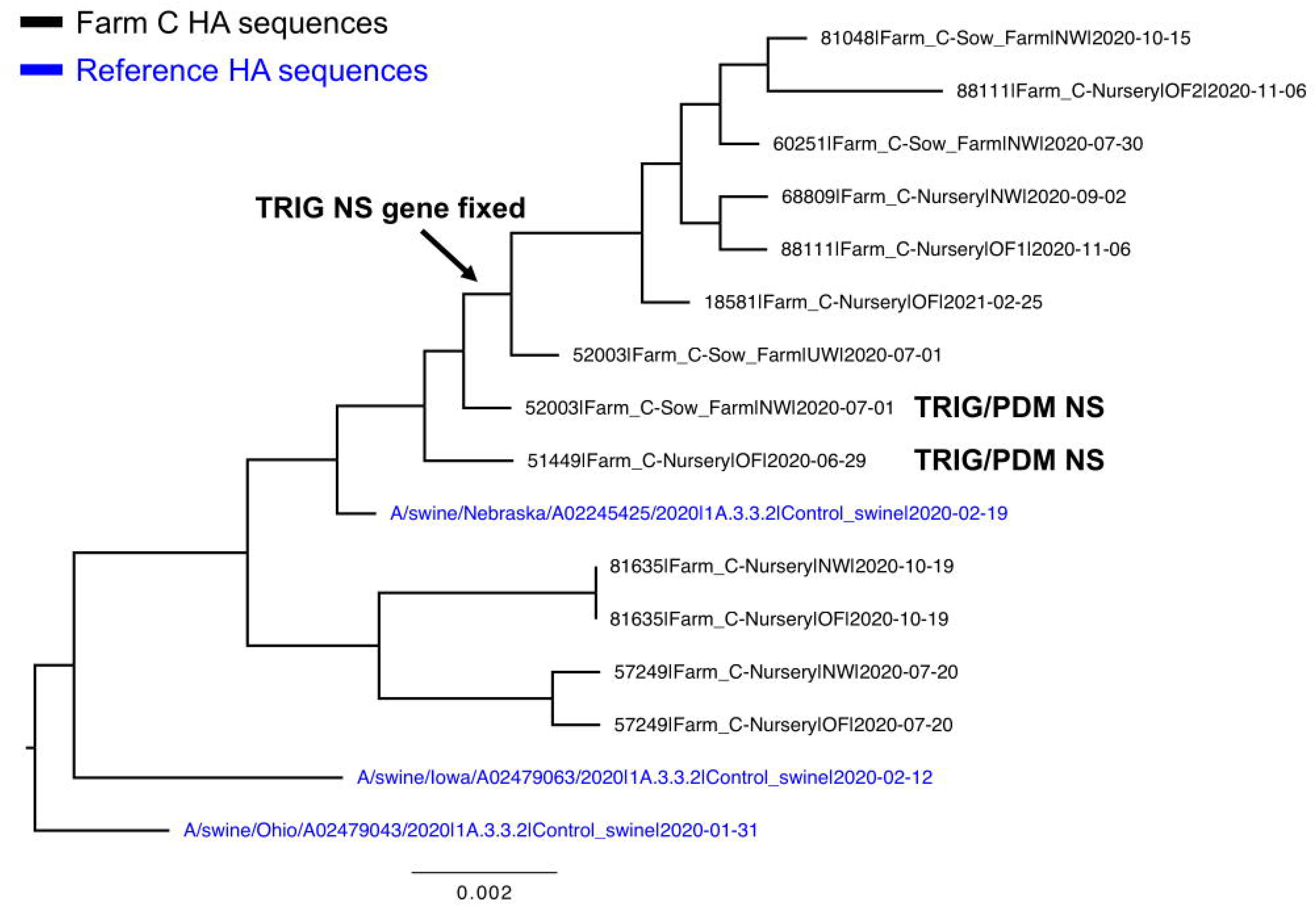
**Reassortment example in Farm C**. Initial co-circulation of the PDM and TRIG NS genes followed by a fixation of the TRIG gene in a clade of 1A.3.3.2 H1 viruses. This clade was a result of a human-to-swine spillover of the 2019-20 season H1N1pdm virus. It captures in real-time the classic dynamic where a novel H1N1pdm virus obtains TRIG internal genes after a spillover into swine.

### Modeling the probability of IAV detection using farm-level parameters

Predictor variables of IAV status of a sample were represented as nodes in a directed acyclic graph (DAG). They included Farm (prod_sys, responses a, b, c, d), whether there was a positive case in a nursery of the same Farm in the previous month (nursery_pos_prev, responses yes, no, NA), and whether there was a positive case in a sow farm of the same Farm in the last month (sow_pos_prev, responses yes, no, NA). These nodes were also linked in the DAG with vaccine use parameters, which were specific to each of the Farms but included whether the location used 1B.2.1 (vacc_1B.2.1), 1B.2.2 (vacc_1B.2.2), 1A.3.3.3 (vacc_1A.3.3.3), or 2010.1 (vacc_2010.1) components in their vaccines (responses yes, no). A final node in the DAG was included and represented the concluding state of positive or negative detection of IAV in the location (detection, responses yes, no). DAGs using these common nodes were generated separately for the sow farms and the nurseries.

The sow farm DAG had the eight common nodes described above and an additional parameter that described sow herd breeding size (sow_breeding_herd_size, continuous variable). Ten arcs described potential causal relationships between these nodes (five from the whitelist). The causal relationships were determined using logically required (whitelist) and logically inconsistent links (blacklist) between nodes in the DAG (Supplemental File 2). The sow farm DAG had an average Markov Blanket size of 2.67, an average neighborhood size of 2.22, and an average branching factor of 1.11. The nursery DAG had the eight common nodes described above and three additional parameters that described piglet mixing at the site (mix_site, responses yes, no), barn (mix_barn, responses yes, no), and room level (mix_room, responses yes, no). There were 13 arcs describing potential causal relationships between these nodes (7 from the whitelist). It had an average Markov Blanket size of 2.91, an average neighborhood size of 2.36, and an average branching factor of 1.18.

The general pattern observed in the sow farm DAG was that the probability of an IAV positive detection was conditionally dependent on whether there was a positive case in the sow farm in the prior month (sow_pos_prev) and 1B.2.2 component containing vaccine use (vacc_1B.2.2). The data suggested that prior detections increase the probability of a given sample being IAV positive. Specifically, for a given sow farm, if there were no prior detections in the preceding month, the probability of a positive detection in the current month was 9.4%. However, if there had been IAV-positive detections within the sow farm in the past month, the probability of a positive detection increased substantially to 44.5%.

In the nursery DAG, the probability of a positive detection was conditionally dependent on whether there was room mixing at the site (mix_room), whether there had been a positive case in a nursery of the same production system in the previous month (nursery_pos_prev), and 1B.2.2 component containing vaccine use (vacc_1B.2.1). Using conditional probability queries, chances of positive detection of IAV increased from 61.9% to 76.6% if the nursery used room mixing and from 63.4% to 79.1% if the nursery used a vaccine containing a 1B.2.1 component. The data suggested that prior detections increase the probability of a given sample being IAV-positive. Specifically, if there were no prior detections in nurseries of the same production system in the preceding month, the probability of a positive detection this month was 47.3%. If there had been IAV-positive detections within nurseries in the past month, the probability of a positive detection increased to 75.7%.

In both the sow farm DAG and the nursery DAG, usage of vaccines with a specific component was associated with an increased probability of positive IAV detection. These results reflect farm-specific usage of different vaccine formulations. In the case of the sow farm, the 1B.2.2 was associated with a 32.7% increased probability of IAV detection. The only farm to use this particular vaccine was Farm A. The Farm A sow samples tested positive for IAV 68.1% of the time, which is a greater percentage than found at Farm B at 60.4% and Farm D at 40.7%. Farm C sow farms had an even greater percentage of positives (75.5%). Therefore, while the use of 1B.2.2 was not directly associated with the highest probability of positive cases across the study, the use of the vaccine does signal that the sample did not originate from Farm B or D, which, consequently, suggests the increased likelihood of a positive detection. Similarly, the 1B.2.1 vaccine, which was found to increase the probability of a positive detection by 15.7%, was used only by Farms A and C. Nurseries from Farms A and C had the highest proportion of IAV-positive samples (78.0% and 89.1%, as opposed to 69.2% at Farm B and 40.9% at Farm D). A plausible explanation for the relationship between the use of 1B.2.1 and/or 1B.2.2 vaccines in a production system and IAV-positive detection probability increases is that they were conflated with some other unmeasured variable(s) that were common to particular subsets of Farms and these factors drove IAV-detection probabilities.

## Discussion

The diversity of IAV in swine in the US has presented a continual challenge for controlling infection in pigs with vaccines [11,12]. Our active surveillance demonstrated that understanding interactions between the different stages of swine production and the IAV diversity within each linked farm is necessary to implement effective vaccine control. Within a single year of surveillance, one production system could harbor three to five HA genetic clades from H1 and H3 subtypes of IAV, even if a single time point only detected one HA genetic clade. A complicating factor was that IAV detection and successful sequencing was impacted by IAV prevalence being lower than 10% in sow farms. We addressed this by developing a statistical framework that supported revising farm-level control based on IAV detections anywhere within a production system. This model was complemented by a phylodynamic analysis showing sow farm-to-nursery transmission routes. Surveillance efforts focusing on sequencing IAV-positive samples from sow farms and maintaining a database of historical IAV sequences and testing results can support control through enhanced biosecurity, quarantine, and/or vaccination. Our data also support the utility of composite or pooled sampling techniques, i.e., oral fluids or udder wipes [58–60], within efficient surveillance protocols. Furthermore, we also reinforced the importance of human-to-swine transmission of IAV [10,41,61]; a dynamic observed across all production systems in both sow and nursery locations. Consequently, biosecurity practices that minimize interspecies transmission will improve herd health and potentially impact global human health.

These data demonstrate how IAV detected within a swine production system reflects the connections between sow farms and nurseries mediated by animal movement. Previous studies have found weaned pigs to be IAV positive upon arrival to the nursery or wean-to-finish sites and have determined the major weaned pig transportation routes correspond with models of IAV dissemination across the U.S. [28,62,63]. The age of first IAV infection is affected by the degree to which piglets are protected by maternally-derived antibodies, which, if conferred, has been suggested to wane over 4-14 weeks [30,64–66]. In our study, we used phylogenetic ancestral state estimation techniques to confirm the occurrence of direct sow farm-to-nursery IAV transmission. In a single production system, Farm D, we could not detect sow farm to nursery transmission patterns for any IAV detections, likely due to low participation and a lack of historical IAV genomic surveillance. Further, in some of our data, viruses detected in the nursery were not recent descendants of those found in the sow farms. The absence of evidence supporting direct transmission routes within a system could be attributed to factors such as physical mixing of weaned piglets from different sow farms, indirect contact with other flows through contaminated spaces such as trucks, aerosolized IAV particles from nearby barns, or human contact leading to interspecies transmission [67–69]. Unsampled sow farm detections cannot be ruled out as a source of these one-off nursery detections. However, our statistical model provides support for system-wide surveillance. Specifically, prior detection of IAV within a system increased the probability of detection in the present, and the HA clades within a system are more likely to circulate in those locations and represent an opportunity to implement control strategies even if premise surveillance cannot detect or sequence IAV. Additionally, though IAV clades are documented to transmit between sow farms and nurseries, our models find case history among farms of the same type (sow farm or nursery) to be the more robust determinant of the probability of a sample testing positive for IAV.

We demonstrate that detecting IAV at the sow farm production level is essential to identify strains that may subsequently be detected in “downstream” production phases, such as in the nursery and finisher pig populations. However, detection and sequencing from sow farm samples can be complicated by low levels of IAV prevalence. Our sampling protocol was designed to detect influenza at 10% prevalence or higher, but recent sow farm active surveillance study has reported a prevalence ranging from 2.0 to 7.5%, and a theoretical modeling study predicts the prevalence of endemic strains to be less than 8% [63,70]. An additional consideration for designing surveillance to improve control strategies is a necessity for consistent sampling efforts. When considering a sow farm and linked nurseries as one unit, we demonstrated 3 to 5 independent HA and NA clades circulating within the unit over a year.

However, at any given time, only a single HA clade could be detected within a premise. Further, our data demonstrated the potential for reassortment within a single system, a phenomenon that can lead to in changes in virus phenotype, consequently altering the detection frequency of specific HA clades within relatively short periods [71,72]. When paired with the amount of HA and genome diversity, this represents a challenge for efficacious autogenous or custom vaccines for a specific location. If one desires to improve animal health and minimize IAV transmission, then we recommend system-level surveillance with a combination of pooled active surveillance samples (NW, UW, OF) and targeted sampling of clinically ill pigs (nasal swabs) to increase the probability of getting the sequence, which could then be utilized for an autogenous vaccine.

When IAV sequences collected before our study were included in analyses, we could document independent nursery detections as recent descendants of viruses endemic to that production system. This further supported the hypothesis that viruses are maintained in sow farms at a low prevalence. Last, the detection of H1N1pdm clade viruses and human-to-swine spillovers in this active surveillance study was higher than expected based on national passive surveillance data. In our study, these spillovers occurred in all four production systems and at both stages of production. We suspect these spillovers commonly remain undetected because they persist sub-clinically [73]. Cross-species spillovers are expected to be self-limiting and rarely maintained [74,75]. However, human-to-swine spillovers occur at relatively high frequencies, and a subset persist for multiple years [10]. Thus, unlike other endemic swine viruses (e.g., porcine reproductive and respiratory syndrome virus), a herd may not remain “IAV negative” unless precautions are consistently taken to reduce the chance of human-to-swine transmission events.

The persistence of multiple genetic clades of endemic swine IAV alongside novel human-to-swine spillovers presents a unique challenge for vaccine control efforts. When human-to-swine spillovers occur, there is potential for reassortment and the formation of novel genotype constellations. This is particularly true in herds where numerous IAV strains co-circulate, as demonstrated in the farms included in this study (Figure 2). The increased genomic diversity likely drives increased HA and NA evolution: reassortment between HA and NA was recently demonstrated to change evolutionary dynamics [21,76] and support the emergence of novel genetic clades of viruses [22]. Genetic changes, when large enough, change the antigenic profile of the virus. If these changes are significant, there is an increased probability that the virus will sweep through the swine herd. As the majority of genes within endemic swine IAV are derived from seasonal human influenzas, it is likely that these antigenically drifted viruses retain the capacity to infect and transmit between humans. The swine H1N1pdm virus clades, in particular, are intricately intertwined phylogenetically with human seasonal pdm viruses, continuously being reintroduced during the human IAV season [9,10]. Previous studies have suggested that IAV is more likely to persist in sow farms because of the constant introduction of new hosts with varying immunologic profiles [77], which provides even greater motivation to include active surveillance at the sow-farm stage of production. Our study suggests that along with improving animal health and minimizing IAV transmission, system-level surveillance is warranted for monitoring reverse zoonoses and detecting IAV with zoonotic potential.

## Data Availability

Datasets and code used in analyses are available at https://github.com/flu-crew/datasets.

## Acknowledgments

We gratefully acknowledge pork producers, swine veterinarians, and laboratories for participating in the USDA Influenza A Virus in Swine Surveillance System and publicly sharing sequences. This work was supported in part by: the Iowa State University Veterinary Diagnostic Laboratory; the U.S. Department of Agriculture (USDA) Agricultural Research Service [ARS project number 5030-32000-231-000-D]; the National Institute of Allergy and Infectious Diseases, National Institutes of Health, Department of Health and Human Services [contract number 75N93021C00015]; the USDA Agricultural Research Service Research Participation Program of the Oak Ridge Institute for Science and Education (ORISE) through an interagency agreement between the U.S. Department of Energy (DOE) and USDA Agricultural Research Service [contract number DOE contract number DE-SC0014664);]; the Department of Defense, Defense Advanced Research Projects Agency, Preventing Emerging Pathogenic Threats program [contract number HR00112020034]; and the SCINet project and the AI Center of Excellence of the USDA Agricultural Research Service (ARS project numbers 0201-88888-003-000D and 0201-88888-002-000D). The funders had no role in study design, data collection and interpretation, or the decision to submit the work for publication. Mention of trade names or commercial products in this article is solely for the purpose of providing specific information and does not imply recommendation or endorsement by the USDA, DOE, ORISE, DARPA, or ISU. USDA is an equal opportunity provider and employer.

## Supplementary Material

**Supplemental File 1.** Survey administered to farm veterinarians at the beginning of the study. Questions are separated into four sections: general information, production and health information, influenza vaccination history, and biosecurity. Veterinarians were allowed to leave additional comments in free text at the end of the survey.

**Supplemental File 2**. Predictor variable whitelists and blacklists used in training of sow farm and nursery Bayesian networks. Whitelisted edges represent prescribed connections within the network, and blacklisted edges represent connections excluded from consideration in the network.

**Supplemental Table 1.** Summary of whole genome sequencing results. Multiple lineages of a sequenced gene are separated by a forward slash. Internal gene lineages are abbreviated (P = H1N1pdm, T = triple reassortant, L = live-attenuated influenza virus vaccine). When a gene segment was unable to be sequenced, it is labeled as “N/A.”

