## Supplemental Table 1 for "Active surveillance for influenza A virus in swine reveals within-farm reassortment and cocirculation of distinct subtypes and genetic clades"

| case | date | farm | type | sample type | ct value | HA | NA | PB2 | PB1 | PA | NP | M | NS |
|---|---|---|---|---|---|---|---|---|---|---|---|---|---|
| 40440 | 5/18/2020 | Farm A | Sow Farm | NW | 29.2 | H1 1B.2.1 | N2 1998 | T | T | T | P | P | T |
| 42185 | 5/26/2020 | Farm A | Nursery | OF | 28.4 | H1 1B.2.1 | N2 1998 | T | T | T | P | P | T |
| 51140 | 6/24/2020 | Farm A | Sow Farm | NW | 28.6 | H1 1B.2.1 | N2 1998 | T | T | T | P | P | T/P |
| 54978 | 7/10/2020 | Farm A | Nursery | NW | 26.9 | H1 1B.2.1 | N2 1998 | T | T | T | P | P | T |
| 54978 | 7/10/2020 | Farm A | Nursery | OF | 29.2 | H1 1B.2.1 | N2 1998 | T | T | T | P | P | T |
| 60447 | 7/30/2020 | Farm A | Nursery | NW | 24.8 | H1 1B.2.1 | N2 1998/N2 2002 | T | T | T | P | P | T |
| 60447 | 7/30/2020 | Farm A | Nursery | OF | 23.4 | H1 1B.2.1 | N2 1998 | T | T | T | P | P | T |
| 66854 | 8/25/2020 | Farm A | Nursery | OF | 26.7 | H1 1A.3.3.2 | N1 pdm | P | P | P | P | P | P |
| 66854 | 8/25/2020 | Farm A | Nursery | OF | 29.0 | N/A | N1 pdm | N/A | N/A | N/A | P | P | P |
| 74399 | 9/22/2020 | Farm A | Nursery | NW | 24.9 | H1 1A.3.3.2 | N1 pdm | P | P | P | P | P | P |
| 74399 | 9/22/2020 | Farm A | Nursery | OF | 28.1 | H1 1A.3.3.2 | N1 pdm | P | P | P | P | P | P |
| 85302 | 10/30/2020 | Farm A | Nursery | NW | 31.2 | H1 1A.3.3.2 | N1 pdm | T | T | T | P | P | T |
| 85302 | 10/30/2020 | Farm A | Nursery | OF | 28.7 | H1 1A.3.3.2 | N1 pdm | T | T | T | P | P | T |
| 07482 | 1/27/2021 | Farm A | Nursery | NW | 24.4 | H1 1B.2.1/H1 1A.3.3.2/H3 1990.4.a | N2 1998 | T | T | T | P | P | T |
| 07482 | 1/27/2021 | Farm A | Nursery | OF | 28.1 | H1 1B.2.1 | N2 1998/N2 2002 | T | T | T | P | P | T |
| 10734 | 2/3/2021 | Farm A | Sow Farm | OF | 27.1 | H1 1A.3.3.2 | N1 pdm | P | P | P | P | P | P |
| 13520 | 2/12/2021 | Farm A | Sow Farm | UW | 27.6 | H1 1A.3.3.2 | N1 pdm | P | P | P | P | P | T/P |
| 26406 | 3/23/2021 | Farm A | Sow Farm | UW | 28.0 | H1 1A.3.3.2 | N1 pdm | N/A | N/A | P | P | P | T/P |
| 26406 | 3/23/2021 | Farm A | Sow Farm | NW | 28.0 | H1 1A.3.3.2 | N1 pdm | P | P | P | P | P | P |
| 34200 | 4/19/2021 | Farm A | Sow Farm | NW | 30.3 | H1 1A.3.3.2 | N1 pdm | N/A | P | P | P | P | P |
| 34200 | 4/19/2021 | Farm A | Sow Farm | UW | 30.4 | H1 1A.3.3.2 | N1 pdm | N/A | P | N/A | P | P | T/P |
| 46559 | 5/25/2021 | Farm A | Nursery | OF | 32.1 | H3 2010.1 | N2 2002 | T | T | T | P | P | T |
| 57423 | 6/30/2021 | Farm A | Nursery | OF | 26.5 | H1 1A.3.3.2/H3 2010.1 | N1 pdm/N2 2002 | T/P | T/P | T/P | P | P | T/P |
| 57423 | 6/30/2021 | Farm A | Nursery | NW | 32.7 | H1 1A.3.3.2/H3 2010.1 | N1 pdm/N2 2002 | T/P | T/P | T/P | P | P | T/P |
| 46595 | 6/10/2020 | Farm B | Sow Farm | UW or NW | 32 or 34.4 | H1 1A.3.3.2 | N1 pdm | T | T | P | P | P | T/P |
| 62522 | 8/7/2020 | Farm B | Nursery | NW | 29.4 | H3 2010.1 | N2 LAIV-98/N2 2002 | T | T | T | T | P | T/P |
| 62522 | 8/7/2020 | Farm B | Nursery | OF | 31.1 | H3 2010.1 | N2 2002 | T | T | T | T | P | T/P |
| 69926 | 9/7/2020 | Farm B | Nursery | NW | 27.8 | H3 1990.4.a | N2 2002 | T | N/A | T | T | P | T |
| 69926 | 9/7/2020 | Farm B | Nursery | OF | 27.0 | H3 1990.4.a | N2 2002 | T | T | T | T | N/A | T/P |
| 69926 | 9/7/2020 | Farm B | Nursery | OF | 26.3 | H3 1990.4.a/H3 2010.1 | N2 2002 | T | T | T | T | P | T |
| 77976 | 10/6/2020 | Farm B | Nursery | OF | 26 | H3 2010.1 | N2 2002 | T | T | T | T | P | T |
| 77976 | 10/6/2020 | Farm B | Nursery | OF | 24.7 | H3 1990.4.a | N2 2002 | T | T | T | T | P | T |
| 88748 | 11/10/2020 | Farm B | Nursery | OF | 31.8 | H3 2010.1 | N2 2002 | T | T | T | T | P | T |
| 88748 | 11/10/2020 | Farm B | Nursery | NW | 28.1 | H3 1990.4.a | N2 2002 | T | N/A | T | T | P | T |
| 95719 | 12/4/2020 | Farm B | Nursery | OF | 27.4 | H1 1A.3.3.3 | N1 classical | N/A | N/A | N/A | P | P | T |
| 95719 | 12/4/2020 | Farm B | Nursery | OF | 22.2 | H1 1A.3.3.3 | N1 classical | T | T | T | P/T | P | T |
| 95719 | 12/4/2020 | Farm B | Nursery | NW | 25.2 | H1 1A.3.3.3 | N1 classical | T | T | T | P | P | T |
| 00315 | 1/5/2021 | Farm B | Nursery | OF | 31.5 | H1 1A.3.3.3 | N1 classical | T | T | T | P | P | T |

| | | | | | | | | | | | | | |
|---|---|---|---|---|---|---|---|---|---|---|---|---|---|
| 10325 | 2/4/2021 | Farm B | Nursery | OF | 26.9 | H1 1A.3.3.3 | N1 classical/N1 pdm/N2 2002 | T | T | T | P | P | T |
| 10325 | 2/4/2021 | Farm B | Nursery | NW | 20.7 | H1 1A.3.3.3/H1 1B.2.2 | N1 classical | T | N/A | T | P/T | P | T |
| 19544 | 3/3/2021 | Farm B | Nursery | OF | 25.6 | N/A | N/A | N/A | N/A | N/A | N/A | P | T |
| 19544 | 3/3/2021 | Farm B | Nursery | NW | 25.7 | H1 1A.3.3.3 | N1 classical | T | T | T | P | P | T/P |
| 28827 | 4/2/2021 | Farm B | Nursery | OF | 28.7 | H1 1A.3.3.3 | N1 classical | T | T | T | P | P | T |
| 38595 | 5/4/2021 | Farm B | Nursery | OF | 26.7 | H1 1A.3.3.3 | N1 classical | T | T | T | P | P | T |
| 49347 | 6/3/2021 | Farm B | Nursery | NW | 23.1 | H1 1A.3.3.3 | N1 classical | T | T | T | P | P | T |
| 49347 | 6/3/2021 | Farm B | Nursery | OF | 26.1 | H1 1B.2.1/H1 1A.3.3.3/H3 2010.1 | N1 classical/N2 1998/N2 2002 | T | T | T | P/T | P | T |
| 52003 | 7/1/2020 | Farm C | Sow Farm | UW | 27 | H1 1A.3.3.2 | N1 pdm | T | T | T | T | P | T |
| 52003 | 7/1/2020 | Farm C | Sow Farm | NW | 31.7 | H1 1A.3.3.2 | N1 pdm | T | T | T | T | P | T/P |
| 51449 | 6/29/2020 | Farm C | Nursery | OF | 31.8 | H1 1A.3.3.2 | N1 pdm | T | T | T | T | P | T/P |
| 60251 | 7/30/2020 | Farm C | Sow Farm | NW | 28.7 | H1 1A.3.3.2 | N1 pdm | T | T | T | T | P | T |
| 57249 | 7/20/2020 | Farm C | Nursery | OF | 32.9 | H1 1A.3.3.2 | N2 2002 | T | T | T | T | P | T |
| 57249 | 7/20/2020 | Farm C | Nursery | NW | 30.0 | H1 1A.3.3.2 | N2 2002 | T | T | T | T | P | T |
| 68809 | 9/2/2020 | Farm C | Nursery | OF | 23 | H3 1990.4.a | N2 2002 | T | T | T | T | P | T |
| 68809 | 9/2/2020 | Farm C | Nursery | NW | 26.4 | H1 1A.3.3.2/H3 1990.4.a | N1 pdm/N2 2002 | T | T | T | T | P | T |
| 81048 | 10/15/2020 | Farm C | Sow Farm | NW | 33 | H1 1A.3.3.2 | N1 pdm | T | T | T | T | P | T |
| 81635 | 10/19/2020 | Farm C | Nursery | OF | 26.2 | H1 1A.3.3.2 | N2 2002 | T | T | T | T | P | T |
| 81635 | 10/19/2020 | Farm C | Nursery | NW | 24.8 | H1 1A.3.3.2 | N1 pdm/N2 2002 | T | T | T | T | P | T |
| 88111 | 11/6/2020 | Farm C | Nursery | OF | 28.2 | H1 1A.3.3.2 | N1 pdm | T | T | T | T | P | T |
| 88111 | 11/6/2020 | Farm C | Nursery | OF | 26.6 | H1 1A.3.3.2 | N1 pdm | T | T | T | T | P | T |
| 02590 | 12/31/2020 | Farm C | Nursery | OF | 25.0 | H1 1B.2.1 | N2 1998 | T | T | T | P | P | T |
| 02590 | 12/31/2020 | Farm C | Nursery | NW | 25.4 | H1 1B.2.1 | N2 1998 | T | T | T | P | P | T |
| 18581 | 2/25/2021 | Farm C | Nursery | OF | 27.0 | H1 1A.3.3.2/H3 1990.4.a | N1 pdm/N2 2002 | T | T | T | P/T | P | T |
| 18581 | 2/25/2021 | Farm C | Nursery | NW | 26.8 | H3 1990.4.a | N2 2002 | T | N/A | N/A | T | P | T |
| 24534 | 3/22/2021 | Farm C | Nursery | OF | 24.7 | H3 1990.4.a | N1 pdm/N2 2002 | T | T | T | T | P | T |
| 24534 | 3/22/2021 | Farm C | Nursery | NW | 26.0 | H3 1990.4.a | N2 2002 | T | T | T | T | P | T |
| 30955 | 4/9/2021 | Farm C | Sow Farm | NW | 28.6 | H3 1990.4.a | N2 2002 | T | T | T | T | P | T |
| 30955 | 4/9/2021 | Farm C | Sow Farm | UW | 28.2 | H3 1990.4.a | N2 2002 | T | T | T | T | P | T |
| 45433 | 5/21/2021 | Farm C | Nursery | OF | 29.5 | N/A | N2 2002 | N/A | N/A | N/A | N/A | N/A | N/A |
| 45433 | 5/21/2021 | Farm C | Nursery | NW | 28.2 | H3 1990.4.a | N2 2002 | T | T | T | T | P | T |
| 63704 | 8/11/2020 | Farm D | Nursery | NW | 27.0 | H1 1A.3.3.2/H3 1990.4.a | N1 pdm/N2 2002 | P | L/T | P | P | P | T/P |
| 63704 | 8/11/2020 | Farm D | Nursery | NW | 27.1 | H3 1990.4.a | N2 2002 | P | T | P | P | P | T |
| 96110 | 12/7/2020 | Farm D | Sow Farm | NW | 24.5 | H1 1A.3.3.2 | N2 pdm | P | L | P | P | P | P |
| 96109 | 12/7/2020 | Farm D | Sow Farm | NW | 24.0 | H1 1A.3.3.2 | N2 pdm | P | L | P | P | P | P |
| 96849 | 12/9/2020 | Farm D | Nursery | OF | 28.8 | N/A | N/A | N/A | N/A | N/A | N/A | P | P |
| 10072 | 2/2/2021 | Farm D | Nursery | OF | 25.0 | H3 2010.1 | N2 2002 | T | L | N/A | P | P | T |
| 38499 | 4/29/2021 | Farm D | Sow Farm | NW | 26.8 | N/A | N2 2002 | T | L | L | N/A | P | T |
| 38499 | 4/29/2021 | Farm D | Sow Farm | UW | 30.2 | N/A | N/A | N/A | N/A | N/A | T | P | T |

| | | | | | | | | | | | | | |
|---|---|---|---|---|---|---|---|---|---|---|---|---|---|
| 37430 | 4/23/2021 | Farm D | Nursery | NW | 30.7 | H3 2010.1 | N2 2002 | T | L | L | P | P | T/P |
| 37430 | 4/23/2021 | Farm D | Nursery | OF | 24.9 | N/A | N/A | N/A | N/A | N/A | N/A | P | T |
