## Supplemental File 1 for "Active surveillance for influenza A virus in swine reveals within-farm reassortment and cocirculation of distinct subtypes and genetic clades"

### Iowa State University Influenza Project 2020-2021
### Farm Questionnaire

**General information:**

1. Sow farm:
    a. Farm name:
    b. State:
2. Herd size
    a. Number of sows (P1-P?) in the breeding herd:
    b. Number replacement gilts (P0) in the breeding herd:
    c. What is the approximate percentage of gilts vs. sows (P1 or higher) in the herd:
3. Nursery farm location
    a. State only:
4. Approximate size of the nursery/nurseries:
    a. Site:
    b. Barn or building:
    c. Room size:
5. Nursery site questions specific to the sampled population during the project:
    a. Is the nursery <u>site</u> single source or mixed:
    b. Is the nursery <u>barn/building</u> single source or mixed:
    c. Is the nursery <u>room</u> single source or mixed:

**Production and health information:**

1. Replacement gilt source
    a. External purchase (yes/no; state):
    b. Internal multiplication (yes/no; state):
2. Replacement gilt location (Isolation; GDU)
    a. Off-site location (yes/no; state):
    b. On-site location (breeding farm yes/no; state):

**Influenza Vaccination History:**

1. Gilt vaccination (yes/no):

    a. Target age at vaccination:
    b. Number of doses prior to entry to breeding herd:
    c. Time of vaccination (# weeks prior to entry to breeding herd):
2. Herd vaccination (yes, no):
    a. Herd vaccination protocol:
        i. When/how often is the herd vaccinated for influenza:
3. Influenza vaccine product:
    a. Commercial or autogenous product:
        i. Name of commercial product:
        ii. Antigen used if autogenous (farm-specific) product:
            1. How many antigens are included in the autogenous product:
            2. What subtypes and specific clusters of H1/H3 in the autogenous product:
            3. What population was the autogenous vaccine virus isolated from:
4. Influenza diagnostics:
    a. Routine diagnostics conducted for influenza:
    b. Post-vaccination diagnostics:

**Biosecurity**

1. What is the quarantine protocol for new gilts:
2. What is the influenza status of the gilt source:
3. Personal protective equipment worn when in contact to pigs:
4. Workers vaccinated for influenza (yes/no):

<u>Additional comments</u>:

We appreciate the time and effort veterinarians have committed to this project and we hope the information will be useful. Again, our **thanks** for your help and assistance with this project.

Phil Gauger
Iowa State University Veterinary Diagnostic Laboratory
