## Supplemental File 2 for "Active surveillance for influenza A virus in swine reveals within-farm reassortment and cocirculation of distinct subtypes and genetic clades"

| from | to | list |
| --- | --- | --- |
| prod_sys | vacc_gamma | nursery_white_list |
| prod_sys | vacc_delta1 | nursery_white_list |
| prod_sys | vacc_delta2 | nursery_white_list |
| prod_sys | vacc_2010_human_like | nursery_white_list |
| prod_sys | mix_site | nursery_white_list |
| prod_sys | mix_room | nursery_white_list |
| prod_sys | mix_barn | nursery_white_list |
| vacc_gamma | vacc_delta1 | nursery_black_list |
| vacc_gamma | vacc_delta2 | nursery_black_list |
| vacc_gamma | vacc_2010_human_like | nursery_black_list |
| vacc_delta1 | vacc_gamma | nursery_black_list |
| vacc_delta1 | vacc_delta2 | nursery_black_list |
| vacc_delta1 | vacc_2010_human_like | nursery_black_list |
| vacc_delta2 | vacc_gamma | nursery_black_list |
| vacc_delta2 | vacc_delta1 | nursery_black_list |
| vacc_delta2 | vacc_2010_human_like | nursery_black_list |
| vacc_2010_human_like | vacc_gamma | nursery_black_list |
| vacc_2010_human_like | vacc_delta1 | nursery_black_list |
| vacc_2010_human_like | vacc_delta2 | nursery_black_list |
| detection | sow_pos_prev | nursery_black_list |
| detection | nursery_pos_prev | nursery_black_list |
| prod_sys | detection | nursery_black_list |
| detection | prod_sys | nursery_black_list |
| prod_sys | vacc_gamma | sow_farm_white_list |
| prod_sys | vacc_delta1 | sow_farm_white_list |
| prod_sys | vacc_delta2 | sow_farm_white_list |
| prod_sys | vacc_2010_human_like | sow_farm_white_list |
| prod_sys | sow_breeding_herd_size | sow_farm_white_list |
| vacc_gamma | sow_breeding_herd_size | sow_farm_black_list |
| vacc_delta1 | sow_breeding_herd_size | sow_farm_black_list |
| vacc_delta2 | sow_breeding_herd_size | sow_farm_black_list |
| vacc_2010_human_like | sow_breeding_herd_size | sow_farm_black_list |
| vacc_gamma | vacc_delta1 | sow_farm_black_list |
| vacc_gamma | vacc_delta2 | sow_farm_black_list |
| vacc_gamma | vacc_2010_human_like | sow_farm_black_list |
| vacc_delta1 | vacc_gamma | sow_farm_black_list |
| vacc_delta1 | vacc_delta2 | sow_farm_black_list |
| vacc_delta1 | vacc_2010_human_like | sow_farm_black_list |
| vacc_delta2 | vacc_gamma | sow_farm_black_list |
| vacc_delta2 | vacc_delta1 | sow_farm_black_list |
| vacc_delta2 | vacc_2010_human_like | sow_farm_black_list |
| vacc_2010_human_like | vacc_gamma | sow_farm_black_list |
| vacc_2010_human_like | vacc_delta1 | sow_farm_black_list |

| | | |
|---|---|---|
| vacc_2010_human_like | vacc_delta2 | sow_farm_black_list |
| detection | sow_pos_prev | sow_farm_black_list |
| detection | nursery_pos_prev | sow_farm_black_list |
| prod_sys | detection | sow_farm_black_list |
| detection | prod_sys | sow_farm_black_list |
